# GraphClust2: annotation and discovery of structured RNAs with scalable and accessible integrative clustering

**DOI:** 10.1101/550335

**Authors:** Milad Miladi, Eteri Sokhoyan, Torsten Houwaart, Steffen Heyne, Fabrizio Costa, Björn Grüning, Rolf Backofen

## Abstract

RNA plays essential roles in all known forms of life. Clustering RNA sequences with common sequence and structure is an essential step towards studying RNA function. With the advent of high-throughput sequencing (HTS) techniques, experimental and genomic data are expanding to complement the predictive methods. However, the existing methods do not effectively utilize and cope with the immense amount of data becoming available.

Hundreds of thousands of non-coding RNAs (ncRNAs) have been detected, however, the annotation of these ncRNAs is lacking behind. Here we present GraphClust2, a comprehensive approach for scalable clustering of RNAs based on sequence and structural similarities. GraphClust2 bridges the gap between HTS and structural RNA analysis, and provides an integrative solution by incorporating diverse experimental and genomic data in an accessible manner via the Galaxy framework. GraphClust2 can efficiently cluster and annotate large datasets of RNAs and supports structure probing data. We demonstrate that the annotation performance of clustering functional RNAs can be considerably improved. Furthermore, an off-the-shelf procedure is introduced for identifying locally conserved structure candidates in long RNAs. We suggest the presence and the sparsity of phylogenetically conserved local structures for a collection of long non-coding RNAs.

By clustering data from two CLIP experiments, we demonstrate the benefits of GraphClust2 for motif discovery under the presence of biological and methodological biases. Finally, we uncover prominent targets of doublestranded RNA binding protein Roquin-1, such as BCOR’s 3 UTR that contains multiple binding stem-loops which are evolutionary conserved.

## 1 Background

High throughput RNA sequencing and computational screens have discovered hundreds of thousands of non-coding RNAs (ncRNAs) with putative cellular functionality [1, 2, 3, 4, 5, 6, 7]. Functional analysis and validation of this vast amount of data demand a reliable and scalable annotation system for the ncRNAs, which is currently still lacking for several reasons. First, it is often challenging to find homologs even for many validated functional ncRNAs as sequence similarities can be very low. Second, the concept of conserved domains, which is quite successfully applied for annotating proteins, is not well-established for ribonucleic acids.

For many ncRNAs and regulatory elements in messenger RNAs (mRNAs), however, it is well known that the secondary structure is better conserved than the sequence, indicating the paramount importance of structure for the functionality. This fact has promoted annotation approaches that try to detect structural homologs in the forms of RNA *families* and *classes* [8]. Members of an RNA family are similar and typically stem from a common ancestor, while RNA classes combine ncRNAs that overlap in function and structure. A prominent example of an RNA class whose members share a common function without a common origin is microRNA. One common approach to detect ncRNA of the same class is to align them first by sequence, then predict and detect functionally conserved structures by applying approaches like RNAalifold [9], RNAz [10], or Evofold [11]. A large portion of ncRNAs from the same RNA class, however, have a sequence identity of less than 70%. In this sequence identity range, sequence-based alignments are not sufficiently accurate [12, 13]. Alternatively, approaches for simultaneous alignment and folding of RNAs such as Foldalign, Dynalign, LocARNA [14, 15, 16] yield better accuracy.

Clusters of ncRNA with a conserved secondary structure are promising candidates for defining RNA families or classes. In order to detect RNA families and classes, Will et al. [17] and Havgaard et al. [14] independently proposed to use the sequence-structure alignment scores between all input sequence pairs to perform hierarchical clustering of putatively functional RNAs. However, their applicability is restricted by the input size, due to the high quartic computational complexity of the alignment calculations over a quadratic number of pairs. Albeit the complexity of similarity computation by pairwise sequence-structure alignment can been reduced to quadratic *O*(*n*^2^) of the sequence length [18], it is still infeasible for most of the practical purposes with several thousand sequence pairs. For the scenarios of this scale, alignment-free approaches such as GraphClust [19] and Nofold [20] propose solutions.

A stochastic context-free grammar (SCFG), also known as covariance model (CM), encodes the sequence and structure features of a family in a probabilistic profile. CM-base approaches have been extensively used, e.g. for discovering homologs of known families [21] or comparing two families [22]. Profile-based methods [20, 23] such as Nofold generally rely on a CM database of known families to annotate and cluster sequences by comparing against the profiles, therefore their applicability for *de novo* family or motif discovery is affected by the characteristics of the already known families and the provided models.

The GraphClust methodology uses a graph kernel approach to integrate both sequence and structure information into high-dimensional sparse feature vectors. These vectors are then rapidly clustered, with a linear-time complexity over the number of sequences, using a locality sensitive hashing technique. While this solved the theoretical problem, the use case guiding the development of the original GraphClust work, here as GraphClust1, was tailored for a user with in-depth experience in RNA bioinformatics that has already the set of processed sequences at hand, and now wants to detect RNA family and classes in this set. However, with the increasing amount of sequencing and genomic data, the tasks of detecting RNA family or classes and motif discovery have been broadened and are becoming a standard as well as appealing tasks for the analysis of high-throughput sequencing (HTS) data.

To answer these demands, here we propose GraphClust2 as a full-fledged solution within the *Galaxy* framework [24]. With the development of GraphClust2, we have materialized the following goals, GraphClust2 is: (i) allowing a smooth and seamless integration of high-throughput experimental data and genomic information; (ii) deployable by the end users less experienced with the field of RNA bioinformatics; (iii) easily expandable for up-and downstream analysis, and allow for enhanced interoperability; (iv) allowing for accessible, reproducible and scalable analysis; and (v) allowing for efficient parallelizations over different platforms. To assist the end users, we have developed auxiliary data processing workflows and integrated alternative prediction tools. The results are presented with intuitive visualizations and information about the clustering.

We show that the proposed solution has an improved clustering quality in the benchmarks. The applicability of GraphClust2 will be shown in some sought-after and prevailing domain scenarios. GraphClust2 supports structure probing data such as from SHAPE and DMS experiments. It will be demonstrated that the structure probing information assists in the clustering procedure and enhances the quality. By clustering ncRNAs from *Arabidopsis thaliana* with genome-wide in vivo DMS-seq data, we demonstrate that the genome-wide probing data can in practice be used for homologous discovery, beyond singleton structure predictions. Furthermore, an off-the-shelf procedure will be introduced to identify locally conserved structure candidates from deep genomic alignments, by starting from a custom genomic locus. By applying this methodology to a couple of well-studied long non-coding RNAs (lncRNAs), we suggest the presence and the sparsity of local structures with highly reliable structural alignments. GraphClust2 can be used as a structure motif-finder to identify the precise structural preferences of RNA binding proteins (RBPs) in cross-linking immunoprecipitation (CLIP) data. By comparing public CLIP data from two double-stranded RBPs SLBP and Roquin-1, we demonstrate the advantage of a scalable approach for discovering structured elements. Under subjective binding preferences of Roquin-1 and the protocol biases, a scaled clustering uncovers structured targets of Roquin-1 that are evolutionary conserved. Finally, we propose BCOR’s mRNA as a prominent binding target of Roquin-1 that contains multiple stem-loop binding elements.

## 2 Materials and Methods

### 2.1 Methods overview

#### The clustering workflow

The GraphClust approach can efficiently cluster thousands of RNA sequences. This is achieved through a workflow with five major steps: (i) pre-processing the input sequences; (ii) secondary structure prediction and graph encoding; (iii) fast linear-time clustering; (iv) cluster alignment and refinement, with an accompanying search with alignment models for extra matches; and finally (v) cluster collection, visualization and annotation. An overview of the workflow is presented in Figure 1.

**Figure 1:**
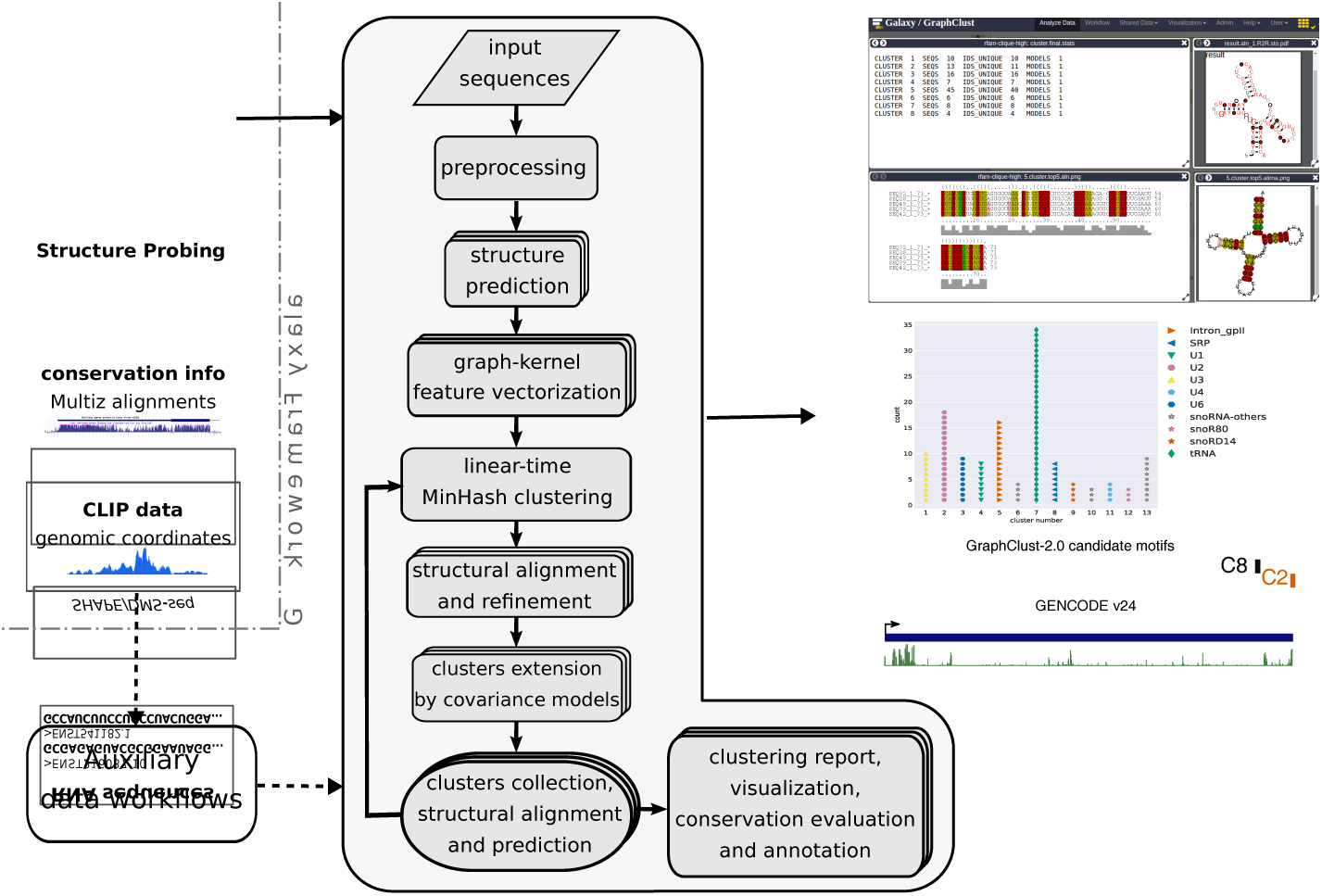
Overview of the GraphClust2 methodology. The flowchart represents the major clustering steps and is supplemented by graphical representations of the associated output data entries. The dashed arrows indicate optional data paths. Auxiliary workflows facilitate integrative clustering of experimental and genomic data including structure probing raw reads or processed reactivities, genomic alignments and conservation information, and genomic intervals e.g. from the CLIP experiments. On the right, a sample selection of the clustering outputs including the overview of the clusters, cluster alignment with LocARNA, RNAalifold consensus structure, and R2R [25] visualization and annotation of the cluster structure by R-scape. Clusters can also be visualised and annotated for the orthology structure conservation predictions.

More precisely, the pre-processed sequences are individually folded according to the thermodynamic free energy models with the structure prediction tools RNAfold [26] or RNAshapes [27]. A decomposition graph kernel is then used to efficiently compute similarity according to the sequence and structure features of secondary structure graphs. The MinHash technique [28] and inverse indexing are used to identify the initial clusters, which correspond to dense neighborhoods of the graph feature space. Formal description and formulations of kernel and MinHash methods are provided in section S1 of the supplementary document. The MinHash clustering approach is very fast with a linear runtime complexity over the number of entries. This accordingly makes GraphClust2 much more efficient than the quadratic all-vs-all approach [19]. It permits the clustering of up to hundreds of thousands of RNA sequences in a reasonable time frame.

After the MinHash clustering step, the *initial* clusters are refined using the RNA domain-specific tools. Firstly, from the sequences of each *initial cluster* a UPGMA tree is created to prune the clusters. The pairwise distances of the tree are approximated from LocARNA sequence-structure alignment scores, as is proposed and detailed in [17]. This pruning procedure keeps the subtree which has the highest average pairwise alignment on its leaves. Here we use the RNA domain-specific scores from LocARNA alignments, although it would have been possible to compute distances from the generic graph kernel scores. LocARNA scores are used, since the runtime complexity is not a concern here, as the pairwise alignments are only computed within each cluster. Each cluster has typically about 10-100 sequences, which is much smaller than the entire input data. In the second step after pruning, the multiple alignment of each pruned cluster is refined with CMfinder’s expectation maximization algorithm [29]. Thirdly, after the alignment is refined, a homology search using Infernal [21] tools is applied over the entire dataset. Such that for each cluster’s refined alignment, a covariance model (CM) is built using cmbuild. The CM is then used to scan the entire sequence database using cmsearch. This CM homology search step extends the clusters with additional homologs that have been missed in the initial clustering. Finally, the sequences of each cluster are aligned with LocARNA, the consensus structures are predicted, visualized and annotated by conservation and covariation metrics.

In an iterative fashion, the steps downstream of the fast clustering can be repeated over the sequences which are not clustered in the previous iteration. GraphClust2 can also compute fuzzy soft overlapping clusters. The option to report overlapping clusters instead of a hard optimal partitioning can be set by the user at the cluster report step. Furthermore, a pre-clustering optional step can be invoked to remove near identical and redundant sequences using CD-HIT [30]. This pre-clustering would be beneficial for the datasets with high redundancy or very large number of sequences, e.g. metatranscriptomics data.

#### Workflow input

GraphClust2 accepts a set of RNA sequences as input. Sequences longer than a defined length are split and processed with a user-defined sliding window option. Two recommended settings are provided for ncRNA clustering and motif discovery as will be discussed in the workflow flavors section below. In addition to the standard FASTA formatted input, a collection of auxiliary workflows are implemented to allow the user to start from genomic coordinate intervals in BED format, or genomic alignments from orthologous regions in MAF format, or sequencing data from the structure probing experiments. Use case scenarios are detailed in the upcoming sections.

#### Workflow output

The core output of the workflow is the set of clustered sequences. Clusters can be chosen either as *hard partitions* having an empty intersection or as overlapping *soft partitions*. In the latter case elements can belong to more than one cluster. In-depth information and comprehensive visualizations about the partitions, cluster alignments and structure conservation metrics are produced (Figure 1). The consensus secondary structure of the cluster is annotated with base-pairing information such as statistically significant covariations that are computed with R-scape [31]. Evaluation metrics for structure conservation are reported. In the case of MAF input, color-coded UCSC tracks are automatically generated to locate and annotate conserved clusters in the genome browser. The in-browser integrated view of the clusterings makes it possible to quickly inspect the results. The Galaxy server keeps track of the input, intermediate and final outputs. The clustering results can be shared or downloaded to the client system.

#### Workflow flavors

Two preconfigured flavors of the workflow are offered for the local and the global scenarios, to facilitate the users without demanding an in-depth knowledge about configuring complex tools. The global flavor aims for clustering RNAs on the whole transcript, such as for annotating ncRNAs of short and medium lengths. The local flavor serves as the motif-finder. The motivation has been to orderly deal with putative genomic sequence contexts around the structured elements. Prediction methods usually require different settings in these two scenarios [32]. The main differences between the flavors are the pre-configured window lengths (*∼*250 vs. *∼*100), the aligner parameters and the hit criteria of the covariance model search (E-value vs. *bit score*). The motif-finder flavor can be for example used to identify cis-regulatory elements, where it is expected to find structured motifs within longer sequences.

As a feature, the fast clustering can be tuned to weigh in sequence-based features. The graph for each entry consists of two disjoint parts. The primary part is the structure graph where the vertices are labeled with the nucleotides while the backbone and paired bases are connected by edges. Besides the primary part, a *path graph* can be included to represent the nucleotide string (option-seq-graph-t). By including the path graphs, sequence-only information would independently contribute to the feature vectors.

### 2.2 Integration of structure probing data

RNA structure probing is an emerging experimental technique for determining the RNA pairing states at nucleotide resolution. Chemical treatment with reagents like SHAPE (selective 2’-hydroxyl acylation analyzed by primer extension) and DMS (dimethyl sulfate) [33, 34] provide one-dimensional reactivity information about the accessibility of nucleotides in an RNA molecule. Structure probing (SP) can considerably improve the secondary structure prediction accuracy of RNAs [35, 36, 37]. SP-assisted computational prediction methods commonly incorporate the probing data by guiding the prediction algorithms via folding constraints and pseudo-energies [38, 39, 26]. Deigan et al. first introduced the position-specific pseudo-energy terms to incorporate the reactivity information alongside the free energy terms of thermodynamic models [40]. The pseudo-energy term for position *i* is defined as: 

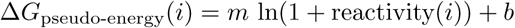

where parameters *m* and *b* determine a scaled conversion of the reactivities to the energy space. GraphClust2 supports structure probing data for enabling a guided structure prediction [26, 41]. The structure probing support is integrated into the pre-processing and the structure prediction steps to generate SP-directed structure graphs.

### 2.3 Implementation and installation

GraphClust2 is implemented within the Galaxy framework [24]. Galaxy offers several advantages to assist our goal of developing a scalable and user-friendly solution. The platform makes it convenient to deploy complex workflows with interoperable tools. Through the uniform user interface across different tools, it is easier for the users to work with new, unfamiliar tools and freely interchange them. Moreover, the standardized data types will ensure that only inputs with valid types are passed to a tool. Interactive tutorial tours are produced to introduce the user interface and guide the user through sample clustering procedures.

GraphClust2’s toolset has been made publicly available in Galaxy ToolShed [42] and can be easily installed into any Galaxy server instance. GraphClust2 is available also as a standalone container solution for a variety of computing platforms at: https://github.com/BackofenLab/GraphClust-2 and can be freely accessed on the European Galaxy server at: https://graphclust.usegalaxy.eu.

#### The workflow implementation

GraphClust2 workflow is composed of tools and scripts which are packaged in Bioconda and Biocontainers [43] and integrated into the Galaxy framework. This has enabled automatic installation of the tools in a version-traceable and reproducible way. All functional units and workflows are manually validated and are under extensive continuous integration tests. Strict versioning of tools and requirements ensures reproducible results over multiple different versions of a tool while delivering updates and enhancements.

#### Platform-independent virtualised container

GraphClust2 can be deployed on any Galaxy server instance, simply by installing the GraphClust tools from the Galaxy ToolShed. As a standalone solution, virtualised Galaxy instance based on Linux containers (Docker, rkt) [44] are provided that can be executed on Linux, OSX and Windows. This largely simplifies the deployment phase, guarantees a reproducible setup and makes it instantiable on numerous computation systems from personal computers to Cloud and high-performance computing (HPC) environments. The Docker image is based on the official Galaxy Docker image [45, 46] and is customized to integrate GraphClust2 tools, workflows and tutorial tours.

### 2.4 Data

#### Rfam-based simulated SHAPE

A set of Rfam [47] sequences and the associated SHAPE reactivities were extracted from the ProbeAlign benchmark dataset [48]. The simulated SHAPE reactivities have been generated according to the probability distributions that are fitted to experimental SHAPE data by Sükösd et al. methodology [49]. Rfam families containing at least ten sequences were used. A uniformly sized subset was also extracted, where exactly ten random sequences were selected per family to obtain a variation with a uniform unbiased contribution from each family.

#### Arabidopsis thaliana ncRNA DMS-seq

*Arabidopsis* DMS-seq reads were obtained from the structure probing experiment by Ding et al. [50] (NCBI SRA entries SRX321617 and SRX320218). The reads were mapped to TAIR-10 ncRNA transcripts (Ensembl release-38) [51]. Reactivities were computed for non-ribosomal RNAs based on the normalized reverse transcription stop counts using Structure-Fold tool in Galaxy [52]. We used Bowtie-2 [53] with the settings recommended by [54] (options –trim5=3, -N=1). Transcripts with poor read coverage tend to bias towards zero-valued reactivities [55]. To mediate this bias, low information content profiles with less than one percent non-zero reactivities were excluded. To focus on secondary structure predictions of the paralogs that can have high sequence similarity, the graphs were encoded with the primary part without path graphs. Information about the ncRNA families is available in the Supplementary Table S4.

#### Orthology sequence extraction from long RNA locus

The genomic coordinates of the longest isoforms were extracted from RefSeq hg38 annotations [56] for FTL mRNA and lncRNAs NEAT1, MALAT1, HOTAIR and XIST. To obtain the orthologous genomic regions in other species, we extracted the genomic alignment blocks in Multiz alignment format (MAF) [57] for each gene using the UCSC table browser [58] (100way-vertebrate, extracted in Aug. 2018). Alignments were directly transferred to the Galaxy server via the UCSC-to-Galaxy data importer. MAF blocks were concatenated using MAF-Galaxy toolset [59] to obtain one sequence per species. An auxiliary workflow for this data extraction procedure is provided. This procedure is notably scalable and can be applied to any locus independent of the annotation availability. Alternatively, the user can provide e.g. full transcripts or synteny regions [60] for the downstream analysis. For the background shuffled input, Multiperm [61] was used to shuffle the Multiz alignment of MALAT1 locus.

#### SLBP eCLIP

Binding sites of SLBP were obtained from the ENCODE eCLIP project (experiment ENCSR483NOP) [62]. In the consortium’s workflow, CLIPper [63] is used to extract peak regions of the read coverage data. The peaks are annotated with both p-values and log2-fold-change scores. These values are determined from the read-counts of the experiment compared with the read-counts of a size matched input. We extracted the peaks with a log2-fold-change of at least 4. To diminish the chance of missing the binding motif, the peak regions were extended by 60 nucleotides both up- and downstream. The sequences of the resulting 3171 binding target regions were used for clustering and motif analysis.

#### Roquin-1 PAR-CLIP

The 16000+ binding sites of Roquin-1 (RC3H1) were obtained from Murakawa et al. [64] (hg18 coordinates from the associated mdc-berlin web page). The 5000 target regions with the highest PAR-CLIP scores were used for the downstream analysis and structural clustering. The binding sites sequences were extracted using the *extract-genomic-dna* tool in Galaxy.

### 2.5 Structure conservation annotation with Evofold, RNAz and R-scape

For each of the studied long RNAs, the sequences were extracted from the orthologous genomic regions as detailed in the data preparation section. Clustering was performed using the motif-finder flavor. In the preprocessing step, the sliding window was set to 100b length and 70% shift. The LocARNA structural alignments of the predicted clusters were further processed using RNAz [65], Evofold [11] and R-scape [31], to *annotate* clusters with structure conservation potentials in the generated genomic browser tracks. RNAz uses a support vector machine (SVM) that is trained on structured RNAs and background to evaluate the thermodynamic stability of sequences folded freely versus constrained by the consensus structure. Evofold uses phylo-SCFGs to evaluate a conservation model for local structures against a competing nonstructural conservation model. R-scape quantifies the statistical significance of base-pair covariations as evidence of structure conservation, under the null hypothesis that alignment column pairs are evolved independently.

RNAz was invoked (option –locarnate) with the default 50% cutoff for SVM-class probability to annotate the clusters. Evofold was also run with the default parameters over the cluster alignments and supplied with the corresponding hg38-100way UCSC’s phylogenetic tree [57]. Clusters that were predicted by Evofold to contain at least one conserved structure with more than three base-pairs were annotated as Evofold hits. R-scape was also applied with the default parameters (i.e. G-test statistics –GTp), clusters with at least two significant covariations were annotated. Clusters are constrained to have a depth of at least 50 sequences. Alignments with spurious consensus structure or no conservation were excluded, using a structure conservation index (SCI) filter of one percent [65]. Clusters annotated by at least one of the three methods are designated as *locally conserved structure candidates*.

### 2.6 Clustering performance metric

The clustering was benchmarked similarly to our previous work [66], such that the Rfam family where each input RNA belongs to is considered as the truth reference class. The performance is measured using the *adjusted Rand index* (ARI) [67] clustering quality metric, which is defined as:

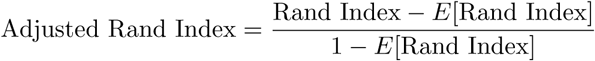

The Rand Index [68] measures the fraction of the entry pairs that are related in the same way in both the predicted clustering and the reference assignment. *E*[Rand Index] is the expected *Rand Index* (for extended details please refer to [66]). The adjusted Rand index is the corrected-for-chance variation of the Rand Index with a maximum value of one. A better agreement between the predicted clustering and the reference assignment leads to a higher ARI value.

## 3 Results and Discussion

### 3.1 Clustering performance evaluation

#### Rfam-cliques benchmark

We evaluated GraphClust2 using known RNA families from the Rfam database [47]. The Rfam sequences were obtained from the *Rfam-cliques* benchmark introduced in our previous work [66]. The Rfam-cliques benchmark contains sets of RNA families at different sequence identity levels and allows for benchmarking a tool for the cases of low and high sequence identities (*Rfam-cliques-low* and *Rfam-cliques-high*). Each variation contains a collection of human members of the Rfam families together with homologs in the other species. As we wanted to evaluate the performance in a simulated scenario of genome-wide screening, we selected the human paralogs from the benchmarks and measured (using the adjusted Rand index metric) how well GraphClust1 and the new pipeline GraphClust2 correctly cluster members of the families together.

In comparison to GraphClust1, GraphClust2 provides alternative approaches for the identification of the secondary structures. Using similar configurations as in GraphClust1 [19], i.e. RNAshapes for structure prediction and bit score for CM search hits, the clustering performance of GraphClust2 is similar or better due to the integration of upgraded tools. However, the alternative configuration of RNAfold for structure prediction and E-value for CM search hits consequently improves the performance (ARI from 0.641 to 0.715 for Rfam-cliques-high, further details in Supplementary Table S1).

#### SHAPE-assisted clustering improves the performance

In the previous benchmark, the clustering relies on the free energy models for secondary structure prediction. A predicted structure sometimes deviates from the real functional structure due to the cellular context and folding dynamics. In this case, the structure probing SHAPE data associated with the real functional structure is expected to improve the quality of structure prediction, which in turn should improve the clustering. We wanted to investigate how an improvement in the structure prediction quality at the early clustering steps influences the final clustering results. To draw a conclusion, however, extensive SHAPE data would be needed for a set of labeled homologous ncRNAs, ideally with different sequence identity and under similar experimental settings. Currently, such collection of data, especially over multiple organisms, is still unavailable. However, as the structure probing is turning into a standard and common procedure, data of such nature is expected to become available soon.

One solution to the mentioned data scarceness is provided in the literature [49], by simulating the experimental generation of a SHAPE profile from the real functional structure. Here, starting from a set of manually curated reference structures, the idea is to simulate SHAPE profiles that reflect the known reference structures. We used the benchmark from ProbeAlign [48] (see Material and Methods for details). Figure 2 shows the effect of incorporating simulated SHAPE data on clustering by guiding the structure prediction. As can be seen, the incorporation of SHAPE data has improved the clustering performance. Notably, an improvement can be achieved in fewer rounds of clustering iterations.

**Figure 2:**
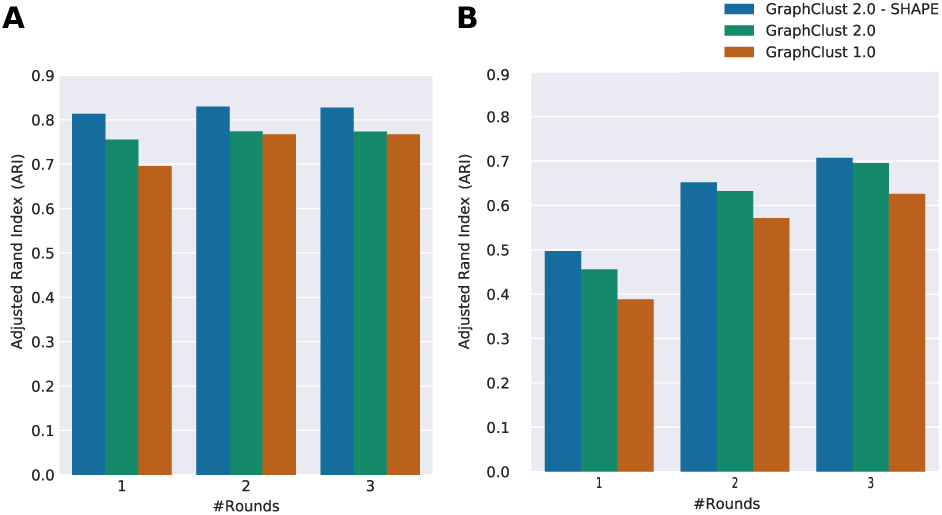
Clustering quality performance over Rfam-based ProbeAlign benchmark dataset and the associated simulated SHAPE data. For comparison, GraphClust2 and GraphClust1 performances are also shown. Incorporating the simulated SHAPE data assists in the clustering performance. (**A**) ARI clustering quality metric for 1-3 rounds of iterative clustering. ARI of the clusterings did not have noticeable improvements after three rounds. (**B**) Similar to (A) but for uniformly sized families, such that precisely ten sequences are randomly extracted per family.

### 3.2 GraphClust2 is scalable

To validate the GraphClust2’s scalability and linearity claims, we used a millions-sequence biological dataset. GraphClust2 is implemented with a comprehensive support for parallel computation using the Galaxy framework. The MinHash-based clustering step is the only step where the entities are evaluated altogether to identify the dense neighborhoods as clusters. Thanks to the MinHash technique, this step has only a linear complexity (see methods and supplementary section S1). To empirically validate this, we clustered a large metatranscriptomic dataset of a marine sample from [70]. After merging the paired-end reads, the metatranscriptome contained 3,594,198 sequences with an average length 250 bases. To filter highly similar sequences, we performed sequence-based pre-clustering with CD-HIT set at a 90% similarity threshold. This produced about 913,000 sequences with a total of 195 million bases. GraphClust2 identified several large clusters of sizes larger than one hundred in one round. Translation-complex-related RNAs (tRNA, LSU and 5s rRNAs) were among the dominating ncRNA classes, matching the expectation due to the high expression levels of the families. Please refer to the supplementary Table S2 for further details. Clustering the entire 3.6 million sequences took less than a day on the European Galaxy server. To check the runtime growth over number of inputs, we measured the wall-clock runtime for sub-samples of various sizes on the European Galaxy server. GraphClust2 robustly scaled with a linear trend over the size of the input (Supplementary Figure S1).

### 3.3 Clustering *Arabidopsis* ncRNAs with DMS-seq in vivo structure probing data

As shown in the previous section, we expect structure probing information to improve the clustering. Information about the structure formations in vivo can be obtained from structure probing (SP) techniques. By determining the nucleotide-resolution base reactivities, where positions with high reactivity indicate unbound bases. Recently, high-throughput sequencing has enabled SP to be applied in a genome-wide manner, thus providing structure probing reactivities of an entire transcriptome [71]. In this way, large amount of SP data can be obtained. Despite the availability of genomic-wide SP data, its application for transcriptome-wide structure analysis is promising [72] but has remained largely underutilized.

#### Enhanced ncRNA annotation with in vivo SP data

We thus evaluated how the task of clustering and annotation of ncRNAs can benefit from such type of genome-wide probing experiments. For this, we compared clusterings of *Arabidopsis thaliana* ncRNAs with and without considering the DMS-seq data by Ding et al. [50] (see Materials and Methods). Due to the relatively high sequence similarity of the annotated paralogous ncRNAs of *Arabidopsis thaliana*, the *Adjusted Rand Index* is high even when no SP data is considered (-DMS-seq mode ARI 0.88). Nonetheless, the quality metric is slightly improved by incorporating the SP data (+DMS-seq mode ARI 0.91). We further manually inspected the quality of the produced clusters. Figure 3 shows the enhanced results for identifying ncRNA classes by using GraphClust2 with in vivo probing data. In the +DMS-seq mode (Figure 3B), all detected clusters are pure RNA classes, while the -DMS-seq mode (Figure 3A) produces mixed-up clusters for *Group II Introns* family plus snoRNA, miRNA and U-snRNA classes. For example, as it can be seen in Figure 3C, the SP data improves the structure prediction by predicting a conserved stem for two of the *Group II Introns* only in the +DMS-seq mode, which leads to one pure cluster for the family.

**Figure 3:**
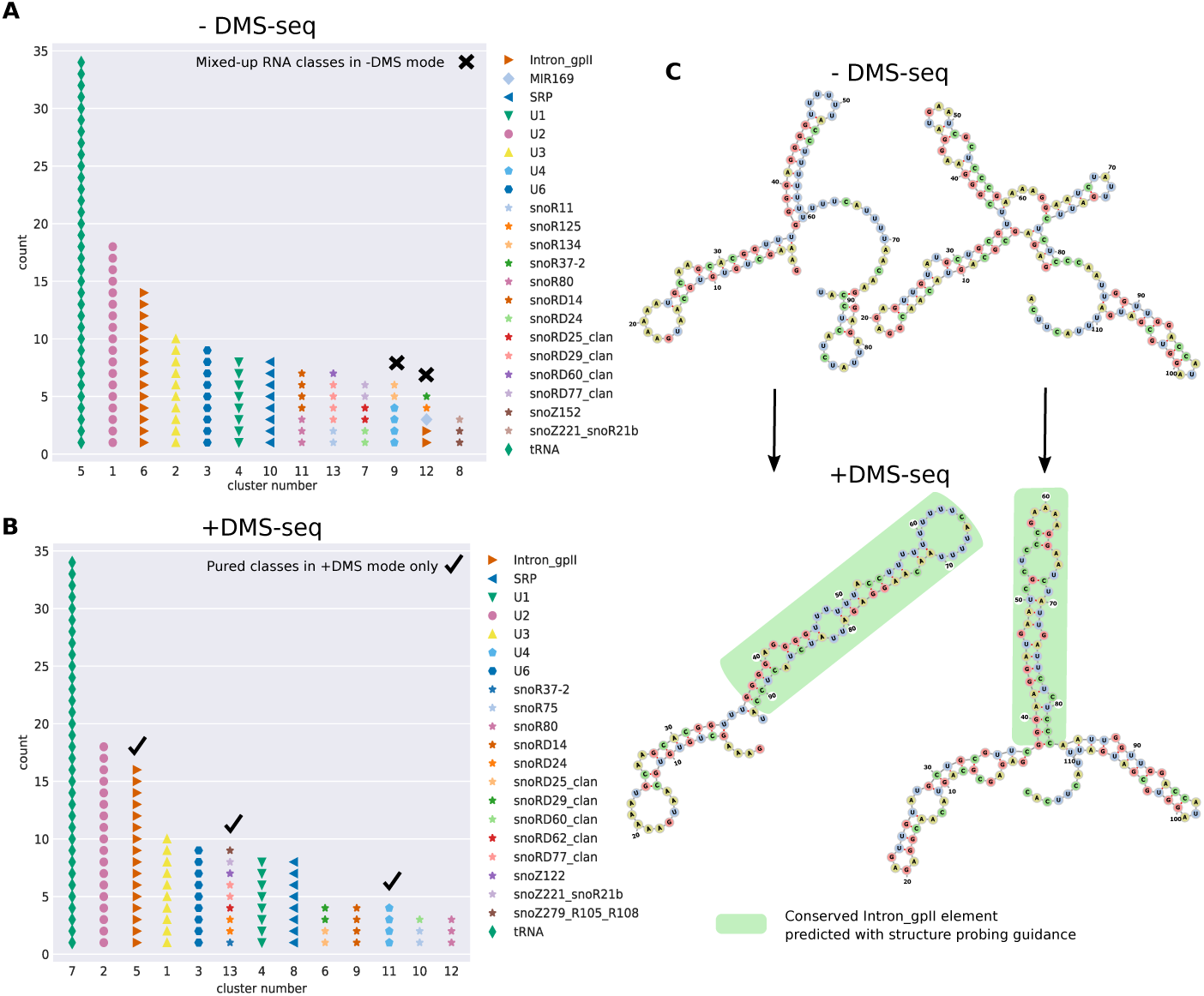
Clustering ncRNAs from *Arabidopsis thaliana* with and without incorporating in vivo DMS-seq structure probing data [50]. (**A**) The predicted clusters without probing data (-DMS-seq) are depicted and the reference family labels are superimposed. Clusters 9 and 12 contain mixed RNA classes. Here, the Group II introns RNA family is split between clusters 6 and 12. (**B**) Similar to A, for +DMS-seq where experimental structure probing data has been used to guide the structure prediction of the generated graphs with pseudo-energy terms. In +DMS-seq mode, only clusters with members from single RNA classes are produced. (**C**) We inspected the predicted structure in more detail. The two transcripts of Intron gpII family are shown that exhibit substantial structure deviations between their MFE structures (-DMS-seq) and the structures guided by the probing data (+DMS-seq). The structures are predicted using Vienna RNAfold and drawn with forna [69]. The highlighted branches correspond to the conserved references structure from Rfam that are correctly predicted only when the DMS-seq reactivities are incorporated.

### 3.4 Discovering locally conserved structured in long RNAs

RNA-seq experiments from biological conditions often result in differentially expressed transcripts, which are studied for functionality and regulatory features. A differential expression hints at putative regulatory effects. An orthogonal source of information for the functional importance of a transcript is phylogenetic conservation patterns. For long non-coding RNAs, however, sequence conservation is usually low, imposing limitations on the sequence level conservation analysis. This fact has been one motivating reason for a collection of recent studies to explore the conservation and functionality of lncRNAs at the secondary structure level [73, 74, 75]. A majority of the studies have been focusing on identifying widely spanned structures, postulating the existence of a to-be-discovered single global structure. However, some of the reported conservations have been challenged for lacking trustworthy base-pair covariations in the alignments [31].

Looking for locally conserved secondary structures in lncRNAs is alluring for several reasons. First, with an increase in the base pair span length the prediction quality decreases [32], which implies that global structure prediction for long RNAs tends to be inaccurate. Second, the structure of a transcribed RNA structure is influenced by RNA-binding proteins in vivo, and thus a predicted global structure likely deviates from the real functional structure. Third, in many cases and similar to the untranslated regions in mRNAs, only a locally conserved structural motif is expected to suffice to perform a function, independent of the precise global structure. We thus revert to a frequently used strategy in the RNA field, namely to look for locally conserved structural motifs. We wanted to evaluate whether we can use GraphClust2 for this purpose.

It should be noted that distinguishing conserved structures from background genomic sequence similarity using base-pair conservation signals is a challenging task. Genome-wide screening studies over genomic alignments require adjusted thresholds for statistical significance discovery and report up to 22% [4] false discovery rates that can be even higher [76]. Despite this and due to the persistent expansion of genomic data, the depth and quality of genomic alignments are continually increasing. Currently, there is a lack of off-the-shelf tools for comprehensively analyzing locally conserved structural elements of a specific locus. Here based on GraphClust2, we propose a data extraction and structure conservation detection methodology (as detailed in the Materials and Methods) that can readily be used for desired loci and genomic alignments to identify *candidates* with locally conserved structure potentials.

An advantage of this clustering approach over traditional screening methods is its ability as an unsupervised learning method, for not imposing explicit presumption on the depth or number of predicted motifs. This makes it possible to find the locally conserved structures also in the regions where a subset of species do not have a conserved structure. Furthermore, this approach does not require a precise co-location of the conserved elements within the transcript, in contrast to traditional alignment-based screening approaches. A further advantage is the availability of the solution in the Galaxy framework, as it provides a rich collection of assets for interactive data collection and analysis of genomic data. We used the 100way vertebrate alignments to extract the orthologous genomic regions for each of the studied RNAs in human and other vertebrates. Each of the orthologous sequences is split into windows, which are then clustered by GraphClust2. The alignment of each cluster has been further annotated with some of best practice complementary methods in assessing covariation patterns and structure conservation potentials, namely RNAz, Evofold and R-scape (see Material and Methods for details). In the following section, some example studies are presented.

### 3.5 Locally conserved candidates with reliable alignments are observable but uncommon

We investigated clustering of orthologous genomic regions of FTL mRNA and four well-studied lncRNAs, using the approach described before. The selected lncRNAs have been previously reported for having loss-of-function phenotypes [77, 78]. In Figures 4A-D and S3 the locations of locally conserved candidates are displayed. These locations are automatically generated by GraphClust2 from clusters with conserved structures (*candidate motifs* track). The track is automatically annotated and filtered using the computed metrics of Evofold, RNAz and R-scape tools (see methods and Figure 4 top-right legend). For these studied lncRNAs, an additional track (*manually curated subset*) is provided. The track is the selection subset of *candidate motifs* track which are manually further screened and selected by stringent expert criteria. The intention was to identify confident conserved elements that can be used e.g. for mutational experiments. The clusters were manually curated in a qualitative manner by inspecting the alignments, their consensus structures as well as the conservation metrics. Only the highly reliable structural alignments which posed a good level of covariation and were not deemed to be alignment artifacts were selected. The main filtering out criteria were: singleton compensatory mutations; avoidable column shifts producing artificial mutations; absence of any region with a basic level of sequence conservation; and similar frequencies of variations in both unpaired and compensatory mutated paired regions. Below we describe the observations from these lncRNAs’ conservation analyses.

**Figure 4:**
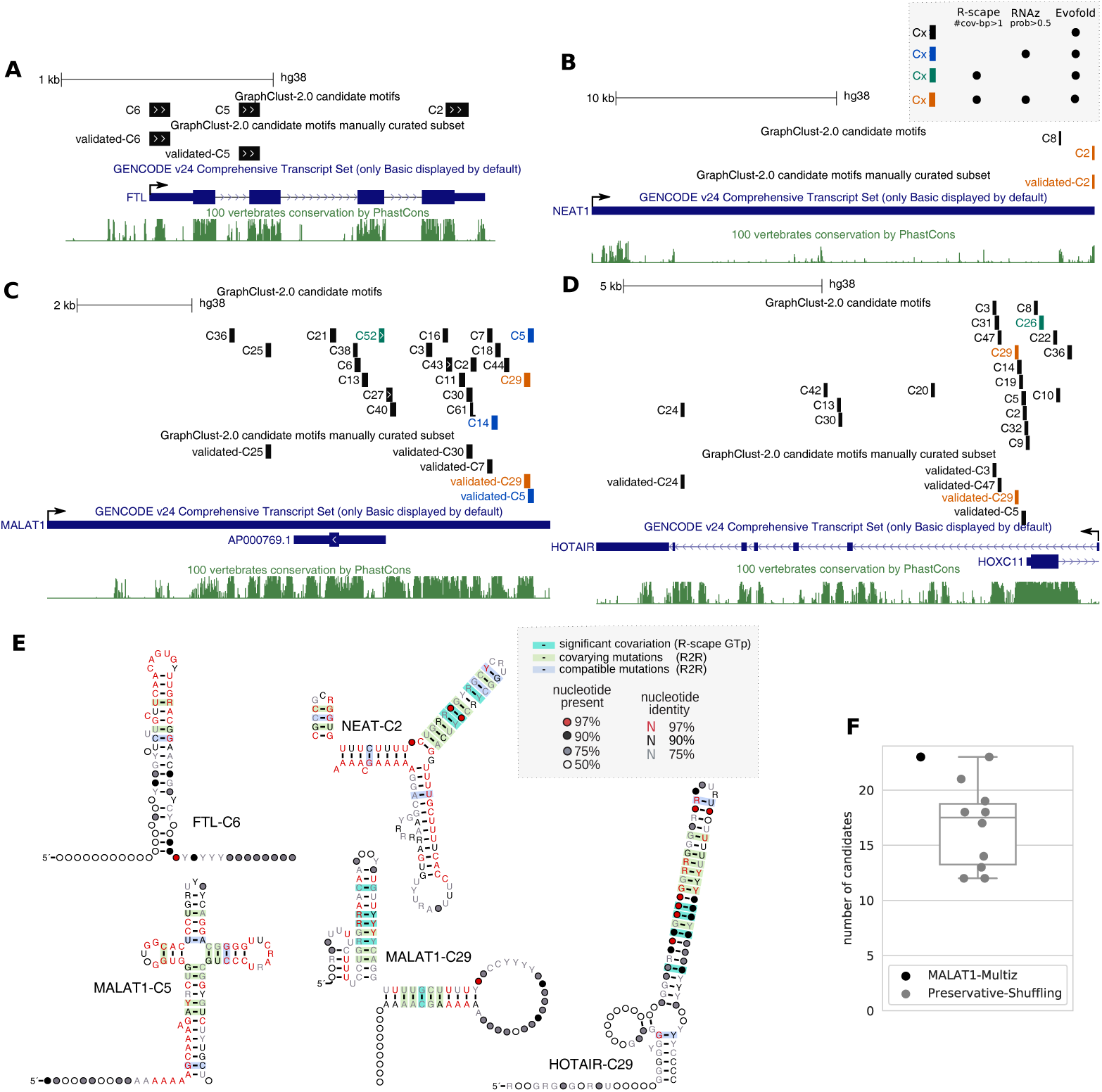
Locally conserved structured elements predicted in FTL mRNA and lncRNAs NEAT1, MALAT1 and HOTAIR. (**A-D**) Locations of the predicted clusters relative to the transcript on the human genome. The clusters under the manually curated subset track, labeled as validated, have passed a qualitative manual screening to exclude unreliable structural alignments (see Results and Discussion). (**E**) Consensus secondary structure for some of the clusters with reliable sequence-structure alignments. Secondary structures are visualized with R2R [25], statistically significant covariation are computed by R-scape and manually overlaid on the R2R visualizations. The alignments are visualized in the Supplementary Figures S6-S11.(**F**) Comparison between the number of predicted candidate motifs of MALAT1 versus ten times Multiperm’s preservative shufflings of the same genomic alignment.

#### FTL

The cis-acting *Iron Response Element* (IRE) is a conserved structured element, that is located on the 5 UTR of *FTL* (*ferritin light chain*) and several other genes. Mutations that disrupt the hairpin structure of IRE cause disease phenotypes by changing the binding affinity of a regulatory Iron Response Protein [79, 80]. As a proof of concept, we applied GraphClust2 to discover structural motifs in the homologous regions of the *FTL* mRNA. The IRE element was identified as one of the three clusters detected by Evofold (Figure 4A).

#### NEAT1

The NEAT1 analysis suggests very limited but also very reliable structure conservation at the 3 end of the transcript that is consensually detected by the three evaluated tools.

#### MALAT1

MALAT1 has relatively a higher level of sequence conservation among the four studied lncRNAs. A higher number of clusters were predicted with a couple of reliable candidates that lean towards the 3 side of the transcript.

To examine how many of the detected motifs are expected to be false positive predictions, we ran the pipeline on ten shufflings of the MALAT1 100way alignment. For the shuffled background, we used Multiperm to preserve the gap structure, local conservation structure patterns and the relative dinucleotide frequencies of the MALAT1 alignment [61]. On average 16.7 candidates were reported for the shuffled genomic alignments, in comparison to the 23 candidates reported for the genomic alignment (Figure 4F). In the predicted candidates set from background, none was commonly annotated by the three methods. For the applied alignment depths and thresholds, Evofold had a considerably higher discovery rate than R-scape and RNAz. In total out of 10 shuffles 167, 9 and 0 clusters were predicted to have a conserved structure by Evofold, R-scape and RNAz respectively.

#### HOTAIR

The predicted candidates for HOTAIR are all located on the intronic regions of the precursor lncRNA. Clustering from the second exon, through skipping the first exon and intron, did not change this observation. A dense number of candidates can be noticed on the first intron that is overlapping with the promoter region of HOXC11 on the opposite strand. Most notably is the candidate cluster HOTAIR-C29, which is highly enriched in G-U wobble base pairs (Figure 4E). In contrast to Watson-Crick GC and AU base pairings, the GU reverse complement AC is not a canonical base pair [81]. Therefore, this structure can only be formed on the antisense RNA and not on the HOXC11’s sense strand.

#### XIST

The XIST candidates are mainly located on the repeat regions and are paralog-like (Figure S3). The manual evaluation of the cluster structural alignments was inconclusive. In the mixture of paralog-like and homolog-like sequences of the cluster alignments, it was not possible to conclude whether the structural variations are merely artifacts of sequence repetition or compensatory mutations of hypothetical structure conservation.

### 3.6 Clustering RNA binding protein target sites

#### SLBP eCLIP

A well-characterized example of an RBP with specific structural preferences is SLBP (*Histone Stem-Loop-Binding Protein*). We clustered target sites of human SLBP using the publicly available eCLIP data [62]. The largest cluster with a defined consensus structure bears statistically significant basepair covariations. The structure matches the SLBP’s Rfam family *“histone 3 UTR stem-loop”* (RF00032). Using the family CM to identify SLBPs on the eCLIP data, we were able to predict exactly the same stem loop structure with the same level of base-pair-covariation (Figure 5A). GraphClust2 and Rfam’s CM hits have more than 95% overlap. These correspondences demonstrate that GraphClust2 can identify the consensus structure element from CLIP data with a high sensitivity.

**Figure 5:**
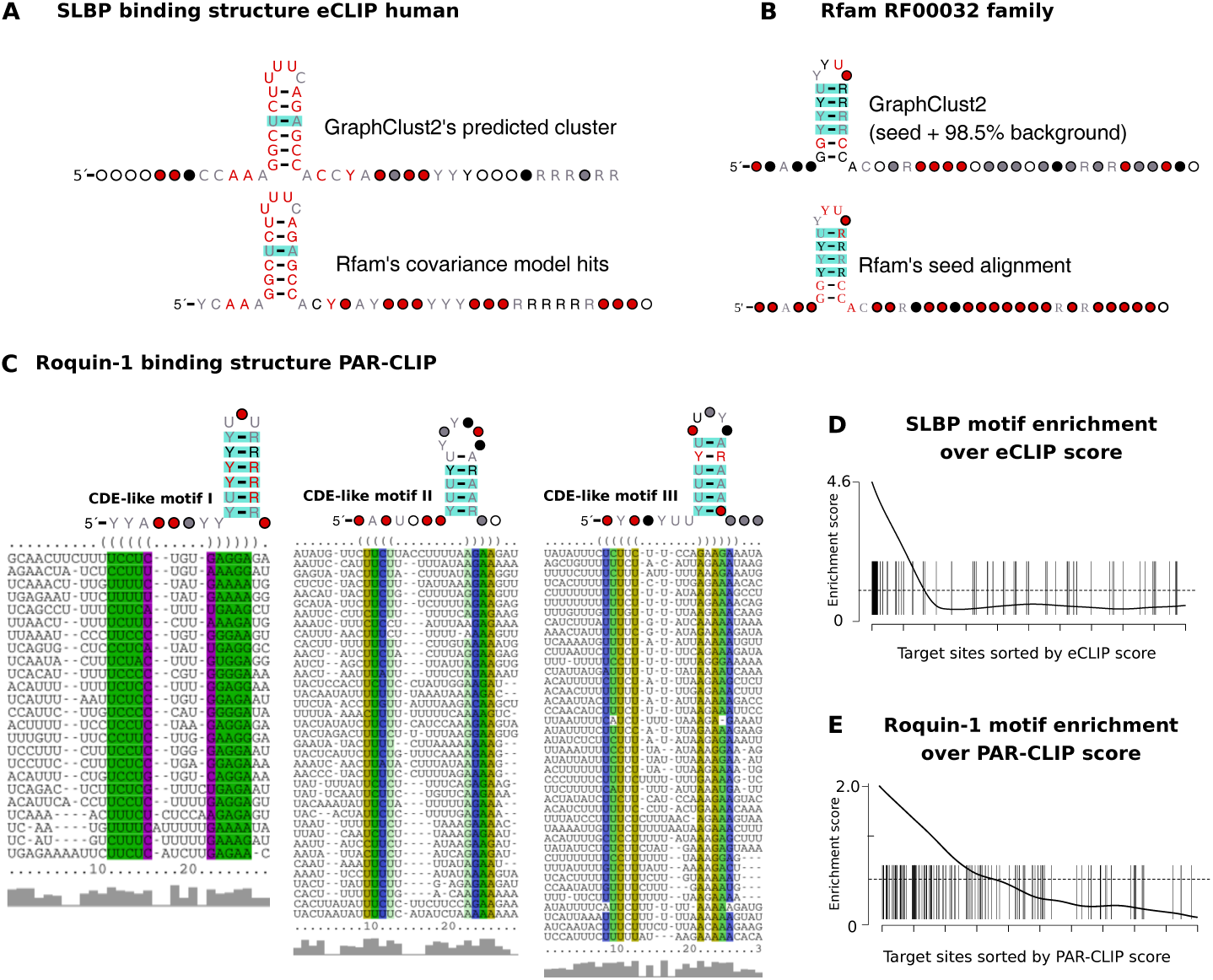
Structured RNA motifs identified by clustering SLBP and Roquin-1 public CLIP data with GraphClust2. (**A**) The consensus secondary structure of the predicted human SLBP motif from eCLIP data versus the consensus structure of cmsearch hits from Rfam’s CM for *histone 3 UTR stem-loop* family RF00032. (**B -top**) The consensus secondary structure of the predicted structure motif from clustering 3000 sequences composed of RF00032’s 46 seed sequences and 2056 background shuffled full sequences as 98.5% background noise. (**B -bottom**) The Rfam’s reference structure for RF00032 seed alignment. (**C**) The consensus secondary structures and alignments of the three clusters with defined consensus structures. The three motifs overlap and have varying loop sizes and uridine content. The structures are akin to the previously validated constitutive decay element (CDE) in TNF-alpha that is a target of Roquin-1. (**C, D**) Gene set enrichment plot of SLBP and Roquin-1 motifs according to the corresponding CLIP scores. SLBP eCLIP has a high enrichment of the stem-loop with strong density in the first hundred target sites. Roquin-1 PAR-CLIP data has a lower enrichment score and low presence in the top 100 target sites. The difference in the enrichment is likely due to the specificity of Roquin-1 that has multiple RNA binding domains and false positive biases of the PAR-CLIP protocol. Scalable clustering assists in overcoming these biases to identify the CDE-like elements. Structures are visualized by R-scape, the color for significant base-pair covariations are adapted to match the legend in Figure 4E. Enrichments are plotted with the Limma R package [82].

The stem-loop structure of the eCLIP data has a lower covariation-level than Rfam’s seed alignment (Figure 5 A vs. B). This is because the Rfam data is phylogenetically diverse (RF00032 seed: 28 species) while eCLIP data is only for Human (eCLIP: K562 cell-line). We checked how GraphClust2 would perform if the eCLIP data from diverse organisms were available. To simulate an eCLIP data with high covariation level, we mixed up the 46 seed sequences of RF00032 family with 2954 shuffled sequences to obtain 3000 sequences such that SLBP is convoluted with 98.5% background. The sequences from the *full* RF00032 set were shuffled to obtain the background of same length and nucleotide content distribution. As can be seen in Figure 5B, GraphClust2 successfully managed to cluster the family entries as one cluster. Here, the cluster has the same stemloop in the consensus secondary structure with the same covariation level as Rfam’s reference structure.

#### Scalable clustering identifies novel CDE-like elements in Roquin-1 PAR-CLIP data

Roquin-1 is a protein with conserved double-stranded RNA binding domains that binds to a constitutive decay element (CDE) in TNF-alpha 3 UTR and several other mRNAs [83, 84]. Roquin-1 promotes mRNA decay and plays an essential role in the post-transcriptional regulation of the immune system [85]. We clustered the binding sites of the publicly available Roquin-1 PAR-CLIP data [64] with GraphClust2. Clustering identified structured elements in three dominant clusters with defined consensus structures. Figure 5C shows the alignments and consensus structures of the three clusters. The consensus structures are similar to the previously reported CDE and CDE-like elements [86].

It should be noted that the union of Roquin-1’s CDE-like motifs have a lower enrichment score based on the PAR-CLIP ranks, in comparison to the SLBP motif based on the eCLIP ranks (Figures 5D,E and S4). For example, only 6 of CDE-like motifs are within the top 100 PAR-CLIP binding sites. Therefore, only the clustering of a broader set of binding targets, with a permissive score threshold, allows identifying the CDE-like elements reliably. We hypothesize that two reasons contribute to the observed distinction. Firstly, eCLIP is an improved protocol with a size-matched input to capture background RNAs of the CLIP protocol [62]. On the other hand, PAR-CLIP is known to have relatively higher false positive rates [87]. Secondly, the ROQ domain of Roquin-1 has two RNA binding sites, one that specifically recognizes CDE-like stem-loops and one that binds to double-stranded RNAs [86, 88]. This would likely broaden the Roquin-1 binding specificity beyond CDE-like stem-loops.

#### BCOR 3 UTR is a prominent conserved target of Roquin-1

We performed a follow-up conservation study over the identified CDE-like motifs from the clustering of Roquin-1 binding sites (Figure 6A). By investigating RNAalifold consensus structure predictions for Multiz alignments of the top 10 binding sites of the conserved candidates, the BCOR’s CDE-like motif was observed to have a highly reliable consensus structure with supporting levels of compensatory mutations. Interestingly the reported CLIP binding site region contains two conserved stem-loops (Figure 6B,C). The shorter stem-loop has a double-sided base-pair covariation and the longer stem-loop contains bulges and compensatory one-sided mutations (Figure 6D,E). Downstream of this site, further binding sites with lower affinities can be seen, where one contains another CDE-like motif. So in total BCOR’s 3 UTR contains three CDE-like motifs (Figure 6F). BCOR has been shown to be a corepressor of BCL6 which is a major sequence-specific transcription repressor. BCL6 expression is tightly regulated and induced by cytokines signaling like Interleukins IL4/7/21 [89, 90]. Overall these results propose BCOR to be a functionally important target of Roquin-1 and assert the role of Roquin-1 in regulating follicular helper T cells differentiation and immune homeostasis pathways [84].

**Figure 6:**
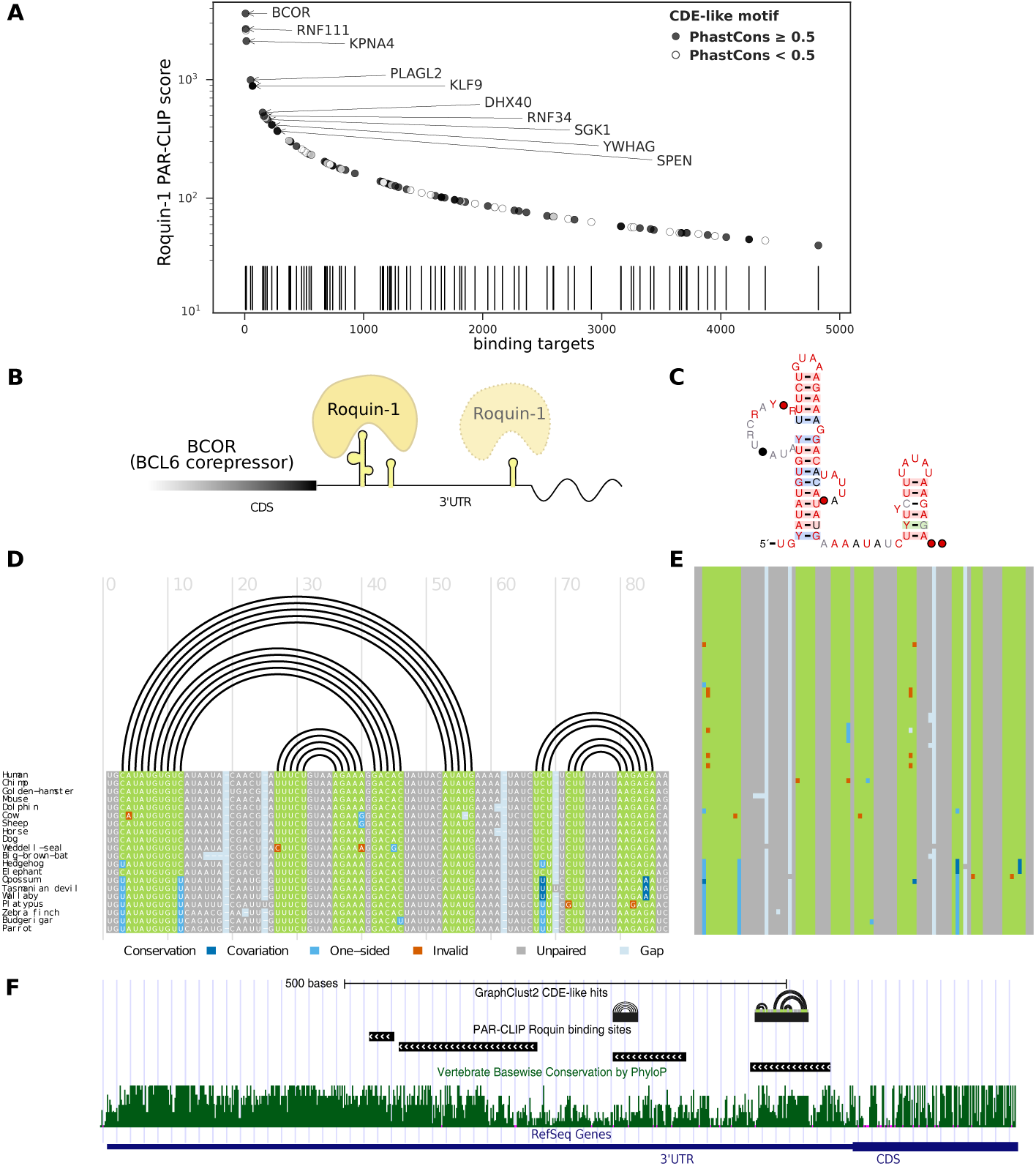
(**A**) The distribution of the Roquin-1 target sites bearing the CDE-like motifs on the 3 UTRs of genes, according to the binding affinity scores. Top 10 highest binding sites with a conserved CDE-like motif are labeled with the associated gene names. (**B**) Roquin-1 binds to a highly conserved double stem-loop element on the 3 UTR of BCOR (BCL6 CoRepressor) with very high affinity. Another CDE-like element with lower affinity downstream of the first element is also spotted. (**C**) R2R visualization for the RNAalifold consensus secondary structure of the conserved double stem-loop element from the vertebrate Multiz alignment. (**D**) Genomic alignment of 20 selected species that is annotated with the consensus structure and the base-pair covariations information. Alignments and compensatory mutations are visualized with R-chie package [91]. (**E**) Genomic alignment overview for the available species extracted from the 100way Multiz alignment. (**F**) Conservation track of BCOR 3 UTR end plus the location of the CDE-like motifs on the negative strand of the locus on the human X chromosome.

## 4 Conclusion

We have presented a method for structural clustering of RNA sequences with a web-based interface within the Galaxy framework. The linear-time alignment-free methodology of GraphClust2, accompanied by cluster refinement and extension using RNA comparative methods and structure probing data, were shown to improve the detection of ncRNA families and structurally conserved elements. We have demonstrated on real-life and complex application scenarios that GraphClust2 provides an accessible and scalable way to perform RNA structure analysis and discovery.

GraphClust2 provides an integrative solution, which can start from raw HTS and genomic data and ends with predicted motifs with extensive visualizations and evaluation metrics. The users can benefit from the vast variety of the bioinformatics tools integrated by the Galaxy community and extend these applications in various ways. Thus, it will be for the first time possible to start from putative ncRNAs in transcriptomic RNA-seq studies and immediately cluster the identified transcripts for annotation purposes in a coherent manner.

## 5 Availability of source code and requirements

- Project name: GraphClust2
- Project repository: https://github.com/BackofenLab/GraphClust-2
- Project home page: https://graphclust.usegalaxy.eu
- Galaxy tools repository: https://github.com/bgruening/galaxytools/tools/GraphClust/
- Operating system(s): Unix (Platform independent with Docker)
- GraphClust2 Docker image: https://hub.docker.com/r/backofenlab/docker-galaxy-graphclust
- License: GNU GPL-v3
- RRID: SCR 017286

## 6 Availability of supporting data and materials

The data presented here that illustrates our work is available from Zenodo [92] and all steps taken for data analysis are accessible via a collection of Galaxy histories from the project homepage at the European Galaxy server (https://graphclust.usegalaxy.eu).

## 7.1 List of abbreviations

ARI: adjusted Rand index;
CDE: constitutive decay element;
CLIP: cross-linking immunoprecipitation;
CM: covariance model;
DMS: dimethyl sulfate;
HPC: high-performance computing;
HTS: high-throughput sequencing;
lncRNA: long non-coding RNA;
MAF: Multiz alignment format;
mRNA: messenger RNA;
ncRNA: non-coding RNA;
RBP: RNA binding protein;
SCFG: stochastic context-free grammar;
SHAPE: selective 2’-hydroxyl acylation analyzed by primer extension;
SP: Structure probing;

## 8 Acknowledgements

We thank Freiburg Galaxy team for their support. We thank Sean Eddy for the helpful comments and discussions. We also thank Sita J. Saunders and Mehmet Tekman for providing feedback about this manuscript.

