## Supplementary Material for "GraphClust2: annotation and discovery of structured RNAs with scalable and accessible integrative clustering"

—

### Supplementary document

Milad Miladi[1], Eteri Sokhoyan[1], Torsten Houwaart[2], Steffen Heyne[3], Fabrizio Costa[4], Björn Grüning[1,5] and Rolf Backofen[1,5,6]

[1]Bioinformatics Group, Department of Computer Science, University of Freiburg, Germany

[2]Institute of Medical Microbiology and Hospital Hygiene, University of Dusseldorf, Germany

[3]Max Planck Institute of Immunobiology and Epigenetics, Freiburg, Germany

[4]Department of Computer Science, University of Exeter, UK

[5]ZBSA Centre for Biological Systems Analysis, University of Freiburg, Germany and

[6]Center for Biological Signaling Studies (BIOSS), University of Freiburg, Germany

#### Contents

**S1 Supplementary methods** **2**
S1.1 NSPDK graph kernel . . . . . . . . . . . . . . . . . . . . . . . . . . 2
S1.2 MinHash technique . . . . . . . . . . . . . . . . . . . . . . . . . . . 2

**S2 Supplementary Tables** **4**

**S3 Supplementary Figures** **6**

**S4 Supplementary tabular files** **15**

### S1 Supplementary methods

#### S1.1 NSPDK graph kernel

Graph kernels can be used to compute the similarity between two graph instances. Here we use the decomposition graph kernel, Neighborhood Subgraph Pairwise Distance Kernel(NSPDK), to evaluate the similarity between the graph-encoded secondary structures. NSPDK defines pairs of subgraphs as neighborhood subgraphs [1, 2]:

**Definition 1.** *For a given graph $G = (V, E)$ and an integer $r \geq 0$ the neighborhood subgraph is a subgraph of $G$ with root vertex $v$ and induced by the set of vertices at distance $d \leq r$. Such subgraph is denoted as $N_r^v(G)$.*

When the distance between the roots of two neighborhood subgraphs of radius $r$ is equal to $d$ the neighborhood-pair relation $R_{r,d}$ holds. Decomposition kernel on that relation $R_{r,d}$ is defined as

$$k_{r,d}(G, G') = \sum_{\substack{A,B \in R_{r,d}^{-1}(G) \\ A',B' \in R_{r,d}^{-1}(G')}} \mathbf{1}(A \cong B')\mathbf{1}(B \cong B') \quad (1)$$

where the inverse relation $R_{r,d}^{-1}$ indicates all possible pairs of neighborhood subgraphs of radius $r$ with root verticies at distance $d$ in the given graph $G$. $\mathbf{1}$ represents the indicator function and $\cong$ stands for isomorphism between the graphs. NSPDK is defined as the sum of all kernels for all radii and all distances.

$$K(G, G') = \sum_{r=0}^{r_{max}} \sum_{d=0}^{d_{max}} k_{r,d}(G, G') \quad (2)$$

An efficient graph serialization procedure is applied to reduce two isomorphic graphs to an identical string to efficiently check for isomorphisms. In the end, an iterative hashing procedure is used to map the string encoding into an integer code. [1] Thus, the isomorphism test between to graphs is reduced to the equality check between their integer codes.

In RNA secondary structure encoded-graphs, for the typically used $r_{max}$ and $d_{max}$ values of range 3-5, the neighborhood-subgraphs would result in sparse features in a high dimensional space. As is detailed in the below section, a local sensitivity hashing scheme is applied to rapidly identify candidate clusters.

#### S1.2 MinHash technique

MinHash [3] is a technique for rapid evaluation of similarity between two sets and also dimensionally reduction. This technique has been successfully applied for large scale clustering and very recently has gained interests from several bioinformatics domains where a rapid evaluation of similarities in large datasets or under high error rates are needed [4]. MinHash is an unbiased estimator with a determinable expected error for the Jaccard similarity coefficient, which is defined as:

$$J(A, B) = \frac{|A \cap B|}{|A \cup B|} \tag{3}$$

, for the two sets $A$ and $B$.

For feature vector $X$ of m dimension $X = \{x_j | 1 \leq i \leq m\}$, and K hash functions $h_i$, the min-hash $hmin_i$ is defined such that:

$$hmin_i = \arg\min_{x_j \in X} h_i(x_j) \tag{4}$$

MinHash function returns the first feature indicator under a random permutation of the features. Using K hash functions for two feature vectors $X$ and $X'$, by comparing their min-hash values we get K estimators of their similarity. If the two instances have $l$ common min-hash values out of the maximum possible $K$, the Jaccard similarity is estimated to be $l/K$.

To obtain an efficient neighbor search procedure, at first all results from the set of MinHash functions are collected in so-called instance sketch to form a signature-like tuple $((hmin_1(X), ..., hmin_K(X)))$. An inverse-index is built for each of K hash-functions, to efficiently with a constant time complexity obtain all the instances that have the same min-hash value. Formally saying, for a given i-th hash function and a value $\bar{h} = hmin_i(X)$, the set of returned instances will be $Z_i(\bar{h}) = \{z \in P | h_i(z) = \bar{h}\}$. Finally, the approximated neighborhood Z is induced from the multi-set $Z = \{Z_i\}_1^N$. In the end, the elements in Z are sorted according to their occurrence frequency. So k-neighborhood of the instance X is the set of k closest elements: $N_k(X)$. Candidate clusters are finally obtained from the densest neighborhoods. The density of the neighborhood for the instance X is defined by the average pairwise similarity between X and all the elements in its k-neighborhood.

### S2 Supplementary Tables

| Dataset | #Rounds | RNAshapes | | RNAfold | |
|---|---|---|---|---|---|
| | | E-val | bitscore | E-val | bitscore |
| Rfam-cliques-low | | | | | |
| | 1 | **0.974** | 0.883 | **0.974** | 0.919 |
| | 2 | **0.977** | 0.887 | 0.975 | 0.922 |
| Rfam-cliques-high | | | | | |
| | 1 | 0.659 | 0.641 | **0.715** | 0.662 |
| | 2 | 0.674 | 0.657 | **0.725** | 0.675 |

Table S1: Clustering performance for the two benchmarking datasets [5] measured by Adjusted Rand Index. Comparison between two alternative methods for generating secondary structure graphs, RNAshapes (version 2.1) and RNAfold (version 2.2), and two Infernal cmsearch hit criteria. GraphClust1 applies RNAshapes with bitscore and GraphClust2 supports all combinations.

| cluster size | size(+pre-clusters) | major class |
|---|---|---|
| 2877 | 6053 | large subunit ribosomal (LSU) rRNA |
| 3395 | 5104 | microRNA mir-598 |
| 535 | 4120 | signal recognition particle (SRP) RNA |
| 347 | 445 | 5S-rRNA |
| 111 | 142 | tRNA |

Table S2: Clustering of exemplary marine metatranscriptome dataset, statistics of the clusters containing non-coding RNAs annotated by Rfam. GraphClust2 identified 28 large clusters of minimum size of 100 in one round of clustering. The second column contain the number of sequences, including the pre-clustered highly similar CD-HIT clusters. Rfam annotations were identified as hit by cmscan against Rfam 14.1 CMs. Only the clusters are listed that were composed of 50% or more ncRNAs.

| experiment | GraphClust2 runtime (hours) |
|---|---|
| NEAT1 | 0.9 |
| MALAT1 | 4.7 |
| HOTAIR | 5.0 |
| XIST | 4.6 |
| FTL | 4.4 |
| Roquin1 | 0.8 |
| SLBP | 0.6 |

Table S3: Runtimes of the local conservation + CLIP experiments on European Galaxy server. It should be noted that the European server is a public resource and the allocation of computation capacity resources is dynamic and depends on the usage load.

| RNA family and clan | #sequences |
|---|---|
| tRNA | 36 |
| U2 | 18 |
| Intron_gpII | 16 |
| U3 | 10 |
| U6 | 9 |
| U1, SRP | 8 |
| U4, snoRD29_clan, snoRD14 | 4 |
| snoRD39_clan, snoR80 | 3 |
| U12, snoZ152, snoU31b , snoRD77_clan, snoRD25_clan, snoRD24, snoR75, snoR134, snoR11 | 2 |
| snoZ43, snoZ279_R105_R108, snoZ221_snoR21b, snoZ199, snoZ159, snoZ155, snoZ122, snoZ105, snoRD74_clan, snoRD62_clan, snoRD60_clan, snoRD44_clan, snoR8a, snoR37-2, snoR135, snoR125, snoR111, MIR414, MIR398, MIR169, MIR163, MIR156 | 1 |

Table S4: Statistics of the ncRNA transcripts extracted from Arabidopsis Thaliana structure probing DMS-seq.

### S3 Supplementary Figures

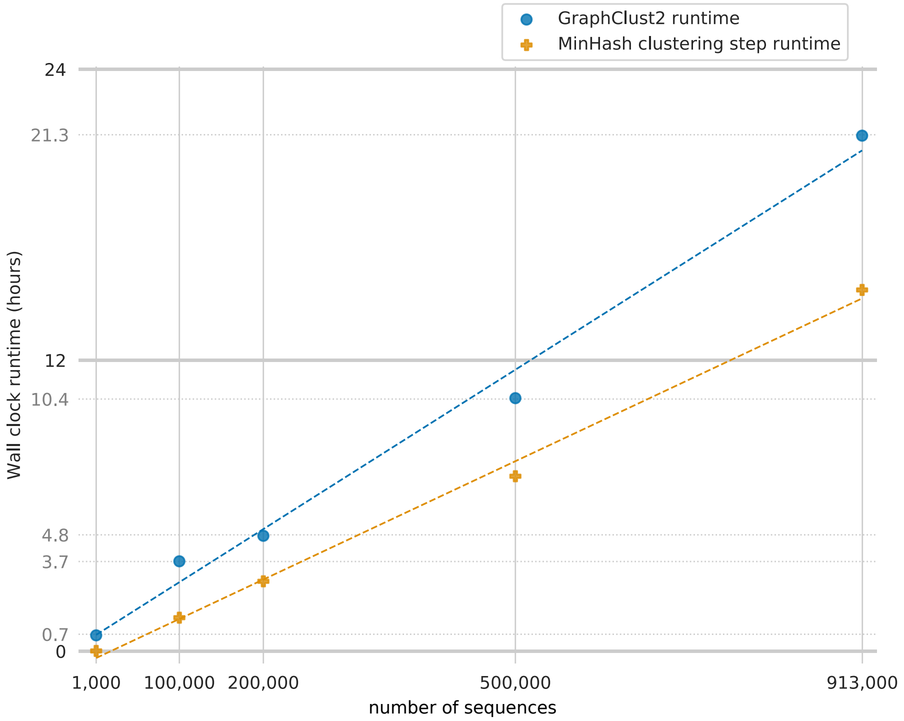


Figure S1: Clustering of exemplary marine metatranscriptome dataset, GraphClust2 runtimes measured on the European Galaxy server. The initial dataset contained 3,594,198 sequences, which were pre-clustered by CD-HIT into 912,675 representative sequences of sequence similarity at most 90%. The 913,000 sequences were iteratively and randomly sub-sampled to obtain the smaller subsets. Using GraphClust2 on the European Galaxy server, each subset was independently clustered in one round, once at a time. It should be noted that the European server is a public resource and the computation capacity resources is dynamically depending on the user load. The linear trend of GraphClust-2 runtime is discernible.

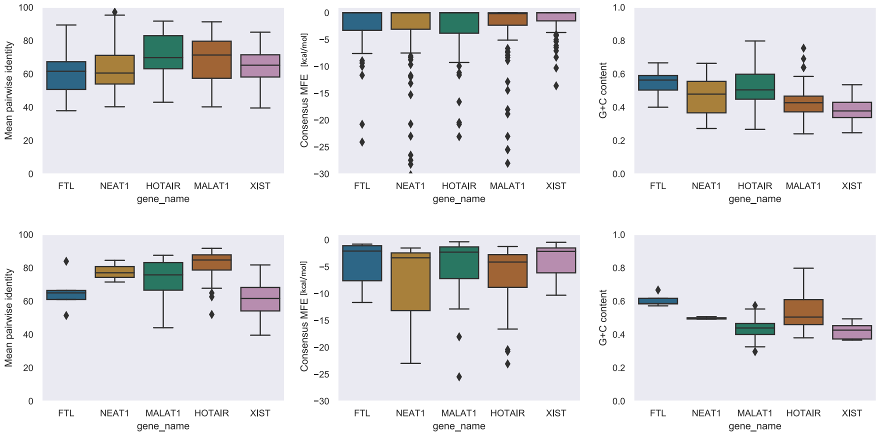


Figure S2: Statistics for the studied long RNAs of all the predicted clusters(Top). For the annotated candidates (Bottom), i.e. the subset of predicted clusters that are annotated with at least one of the three conservation analysis methods (Evofold2, RNAz and R-scape), that are shown in Figures 4 and S3.

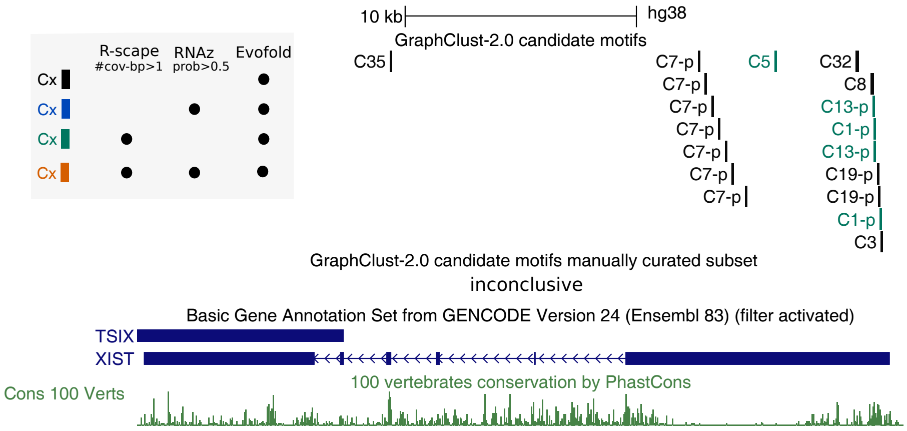


Figure S3: Locally conserved structured elements in XIST lncRNA. Sequences are obtained from 100way vertebrates genomic alignments. Clusters with structure alignments of depth at least 50 and passing one of the three integrated conservation analysis methods. Location of the human sequence of candidate clusters of XIST on the human genome. Paralog-like candidates where multiple human sequences exist in one cluster are suffixed with -p. It must be noted that Evofold predictions are not reliable since the tool is not designed to detect paralog conservation.

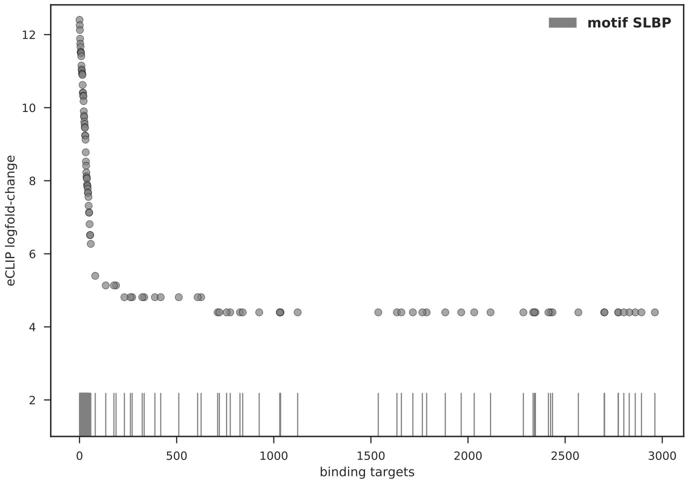


Figure S4: The distribution of predicted SLBP motifs from clustering eCLIP data over eCLIP binding scores. The motif is strongly enriched in the top 100 binding sites.

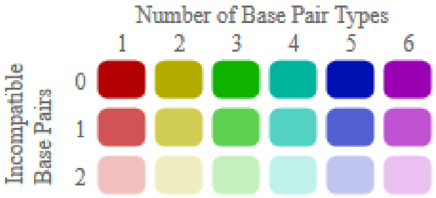


Figure S5: Color legend for LocARNA and RNAalifold alignment visualizations in Figures S6-S11 below. “Compatible base pairs are colored, where the hue shows the number of different types C-G, G-C, A-U, U-A, G-U or U-G of compatible base pairs in the corresponding columns. In this way the hue shows sequence conservation of the base pair. The saturation decreases with the number of incompatible base pairs. Thus, it indicates the structural conservation of the base pair.“ [6]

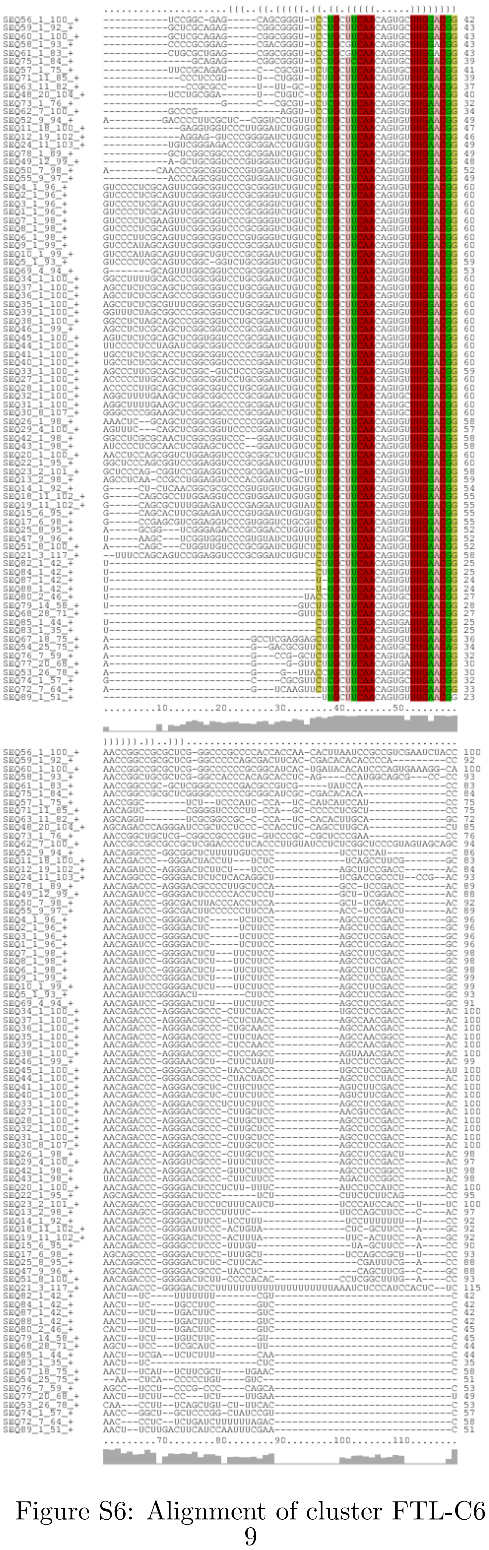

Figure S6: Alignment of cluster FTL-C6

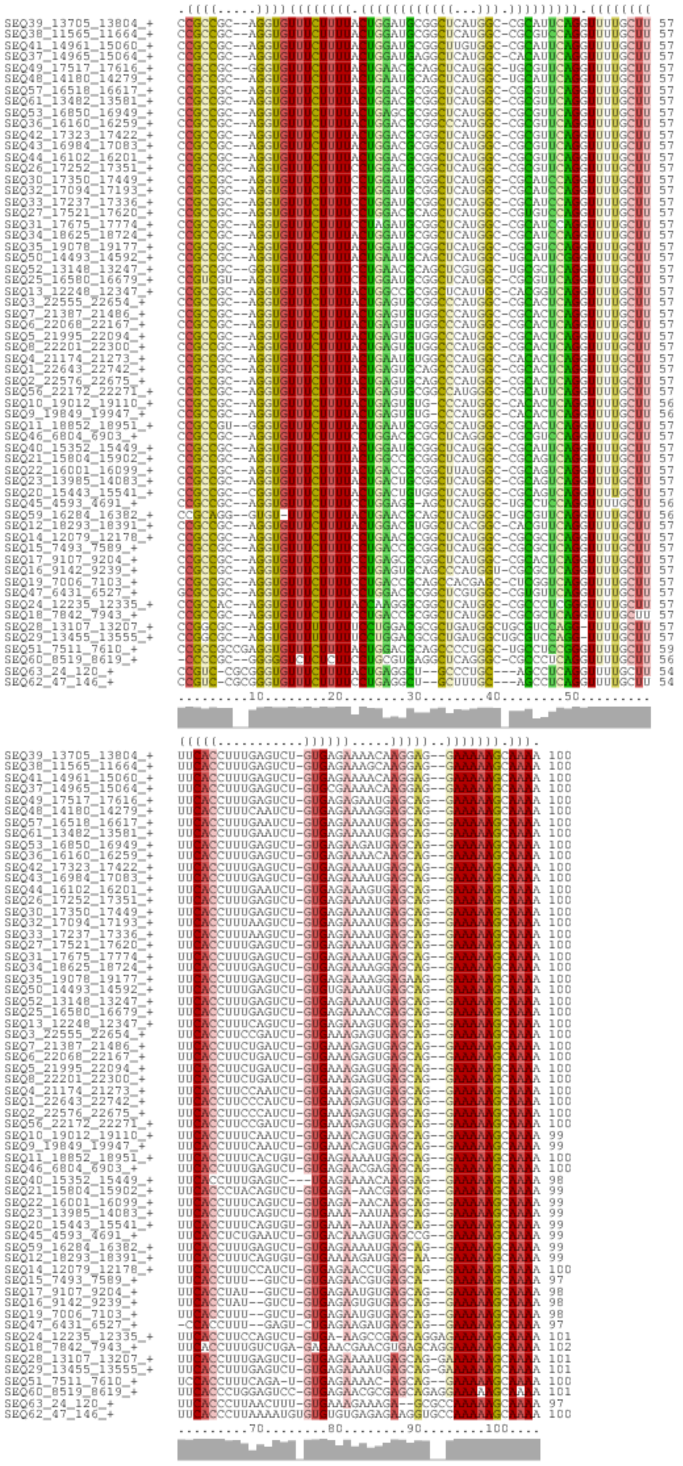

Figure S7: Alignment of cluster NEAT1-C2

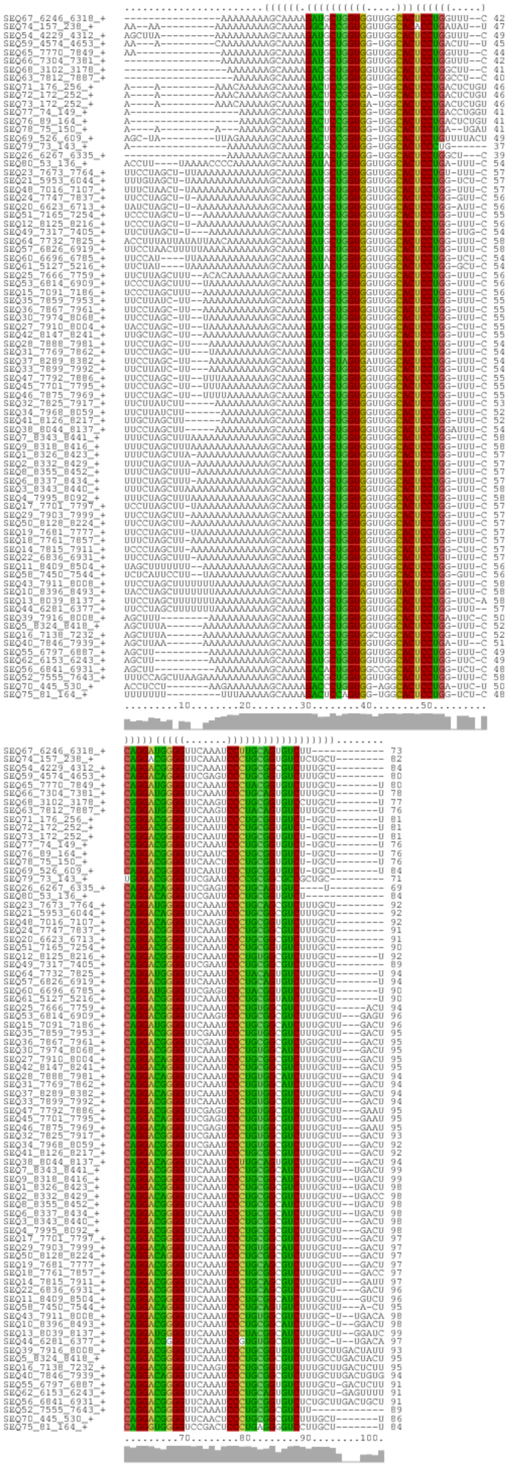

Figure S8: Alignment of cluster MALAT1-C5

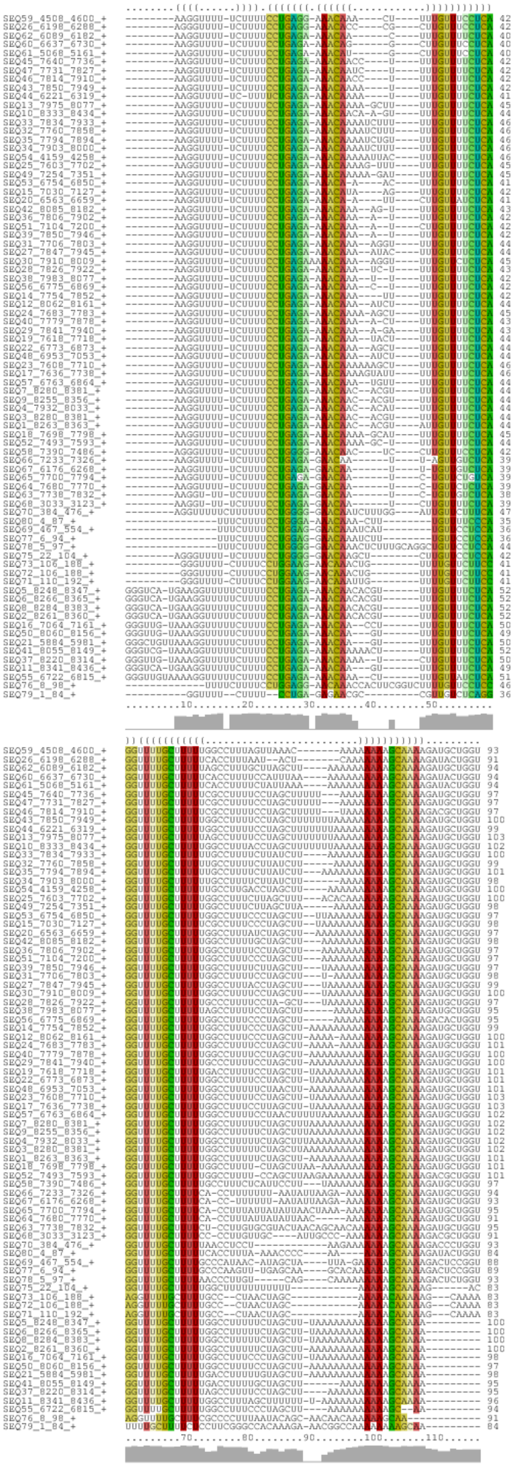

Figure S9: Alignment of cluster MALAT1-C29

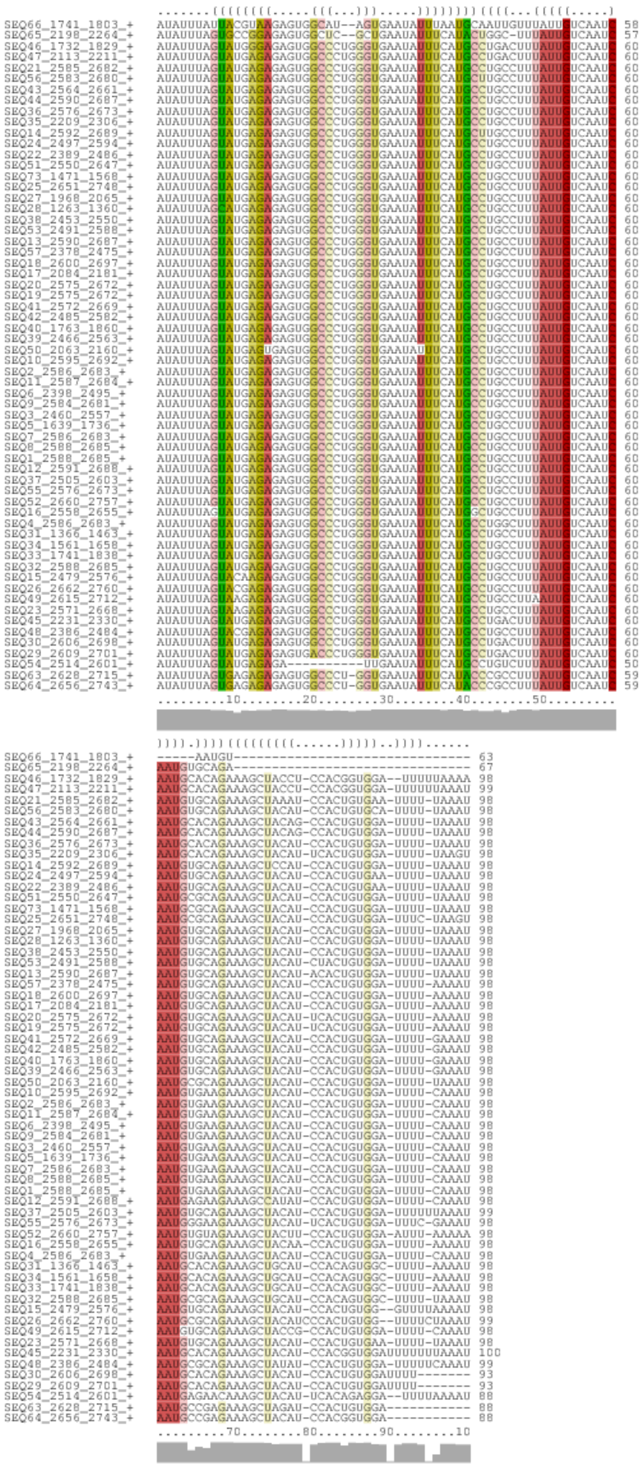

Figure S10: Alignment of cluster HOTAIR-C3

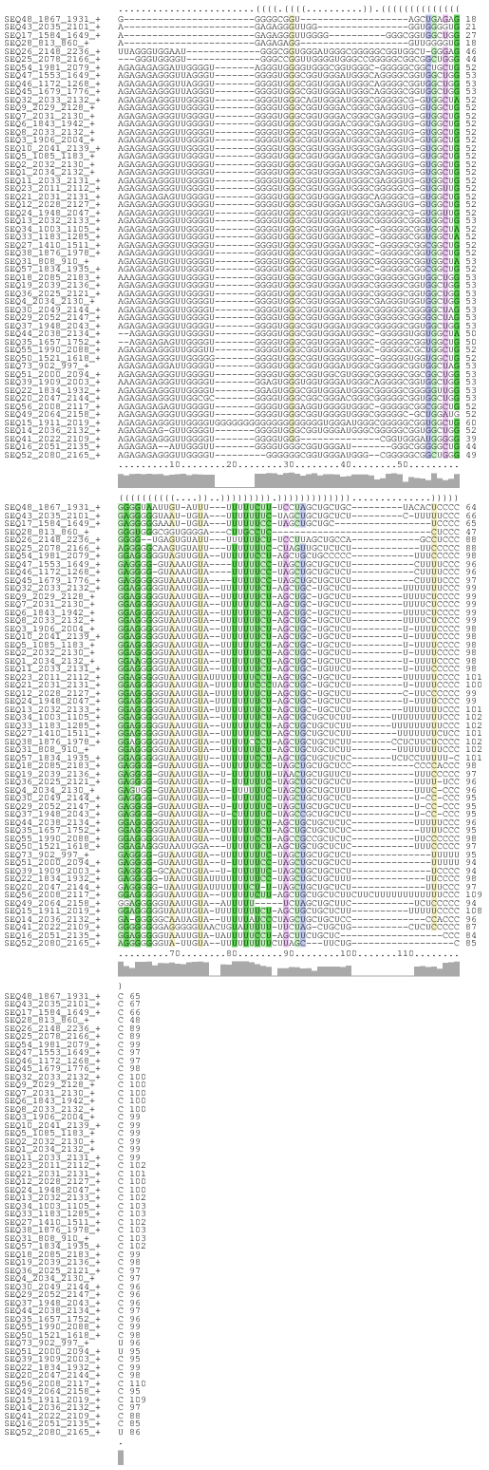

Figure S11: Alignment of cluster HOTAIR-C29

#### S4 Supplementary tabular files

- Tabular file T1: Structure conservation metrics, statistics, and genomic coordinates for the candidates from the long RNA orthologous analysis, corresponding to Figure 4.
- Tabular file T2: Gene names, stem-loop coordinates, conservation info of the CDE-like motifs in 3'UTR by clustering Roquin-1 PAR-CLIP data [7], corresponding to Figures 5 and 6.

[6] Martin Raden, Syed M Ali, Omer S Alkhnbashi, Anke Busch, Fabrizio Costa, Jason A Davis, Florian Eggenhofer, Rick Gelhausen, Jens Georg, Steffen Heyne, Michael Hiller, Kousik Kundu, Robert Kleinkauf, Steffen C Lott, Mostafa M Mohamed, Alexander Mattheis, Milad Miladi, Andreas S Richter, Sebastian Will, Joachim Wolff, Patrick R Wright, and Rolf Backofen. Freiburg RNA tools: a central online resource for RNA-focused research and teaching. *Nucleic Acids Research*, 46(W1):W25–W29, 2018.

[7] Yasuhiro Murakawa, Michael Hinz, Janina Mothes, Anja Schuetz, Michael Uhl, Emanuel Wyler, Tomoharu Yasuda, Guido Mastrobuoni, Caroline C Friedel, Lars Dölken, et al. Rc3h1 post-transcriptionally regulates a20 mrna and modulates the activity of the ikk/nf-$\kappa$b pathway. *Nature communications*, 6:7367, 2015.
